# The opportunistic pathogen *Stenotrophomonas maltophilia* utilizes a type IV secretion system for interbacterial killing

**DOI:** 10.1101/557322

**Authors:** Ethel Bayer-Santos, William Cenens, Bruno Yasui Matsuyama, Giancarlo Di Sessa, Izabel Del Valle Mininel, Chuck Shaker Farah

## Abstract

Bacterial type IV secretion systems (T4SS) are a highly diversified but evolutionarily related family of macromolecule transporters that can secrete proteins and DNA into the extracellular medium or into target cells. They have been long known to play a fundamental role in bacterial conjugation and virulence of several species. It was recently shown that a subtype of T4SS harboured by the plant pathogenic bacterium *Xanthomonas citri* transfers toxins into other bacteria cells resulting in cell death. In this study, we show that a similar T4SS from the multi-drug-resistant global opportunistic pathogen *Stenotrophomonas maltophilia* is proficient in killing competitor bacterial species. T4SS-dependent duelling between *S. maltophilia* and *X. citri* was observed by time-lapse fluorescence microscopy. A bioinformatic search of the *S. maltophilia* K279a genome for proteins containing a C-terminal domain (XVIPCD) conserved in *X. citri* T4SS effectors identified eleven putative effectors secreted by the *S. maltophilia* T4SS. Six of these effectors have no recognizable domain except for the XVIPCD. We selected one of these new effectors (Smlt3024) and its cognate inhibitor (Smlt3025) for further characterization and confirmed that Smlt3024 is indeed secreted in a T4SS-dependent manner by *S. maltophilia* when in contact with a target bacterial species. Expression of Smlt3024 in the periplasm of *E. coli* resulted in greatly reduced growth rate and cell size, which could be counteracted by co-expression with its cognate periplasmic inhibitor, Smlt3025. This work expands our current knowledge about the diverse function of T4SSs subtypes and increases the panel of effectors known to be involved in T4SS-mediated interbacterial competition. Further elucidation of the mechanism of these antibacterial proteins could lead to the discovery of new antibacterial targets. The study also adds information about the molecular mechanisms possibly contributing to the establishment of *S. maltophilia* in different biotic and abiotic surfaces in both clinical and environmental settings.

**Author Summary:** Competition between microorganisms for nutrients and space determines which species will emerge and dominate or be eradicated from a specific habitat. Bacteria use a series of mechanisms to kill or prevent multiplication of competitor species. Recently, it was reported that a subtype of type IV secretion system (T4SS) works as a weapon to kill competitor bacterial species. In this study, we show that an important human opportunistic pathogen, *Stenotrophomonas maltophilia*, harbours a T4SS that promotes killing of competitor species. We also identified a series of new toxic proteins secreted by *S. maltophilia* via its T4SS to poison competitor species. We showed that two different bacterial species that harbour a bacteria-killing T4SS can kill each other; most likely due to differences in effector-immunity protein pairs. This work expands our current knowledge about the bacterial arsenal used in competitions with other species and expands the repertoire of antibacterial ammunition fired by T4SSs. In addition, the work contributes with knowledge on the possible mechanisms used by *S. maltophilia* to establish communities in different biotic and abiotic surfaces in both clinical and environmental settings.

## Introduction

The ecological interactions between bacterial species range from cooperative to competitive and can be mediated by diffusible soluble factors secreted into the extracellular medium or by factors transferred directly into target cells in a contact-dependent manner [1]. Several types of contact-dependent antagonistic interactions between bacteria have been described. Contact-dependent growth inhibition (CDI) is mediated by the CdiA/CdiB family of two-partner secretion proteins composed of the CdiB outer membrane protein that is required for secretion of CdiA, which contains a C-terminal toxin domain [2, 3]. The type VI secretion system (T6SS) is a dynamic contractile organelle evolutionarily related to bacteriophage tails that is attached to the cell envelope, enabling the injection of proteinaceous effectors into target prokaryotic or eukaryotic cells [4, 5]. More recently, contact-dependent antagonism was reported to be mediated via a specialized type IV secretion system (T4SS) that transports toxic effectors into target prokaryotic cell [6].

T4SSs are a highly diverse superfamily of secretion systems found in many species of Gram-negative and Gram-positive bacteria. These systems mediate a wide range of events from transfer of DNA during bacterial conjugation to transfer of effector proteins into infected eukaryotic host cells [7] and into competitor bacteria [6]. T4SSs have been classified based on their physiological functions as (i) conjugation systems, (ii) effector translocators, or (iii) contact-independent DNA/protein exchange systems [8]. Another common classification scheme divides T4SSs into two phylogenetic families called types A and B [9, 10]; while more finely discriminating phylogenetic analyses based on two highly conserved T4SS ATPases (VirB4 and VirD4) identified eight distinct clades [11, 12].

The model type A VirB/D4 T4SS from *Agrobacterium tumefaciens*, which is used to transfer tumour-inducing effectors into some plants species [13], is composed of a core set of 12 proteins designated VirB1-VirB11 and VirD4. Electron microscopy studies on homologous systems from the conjugative plasmids R388 and pKM101 have revealed an architecture that can be divided into two large subcomplexes: i) a periplasmatic core complex made up of 14 repeats of VirB7, VirB9 and VirB10 subunits that forms a pore in the outer membrane and which is also linked, via VirB10, to the inner membrane and ii) an inner membrane complex composed of VirB3, VirB6 and VirB8 and three ATPases (VirB4, VirB11 and VirD4) that energize the system during pilus formation and substrate transfer. Finally, VirB2 and VirB5 form the extracellular pilus and VirB1 is a periplasmic transglycosidase [14-16]. The *X. citri* T4SS involved in bacterial killing shares many features with the type A T4SSs from *A. tumefaciens* and the conjugative T4SSs pKM101 and R388, with one distinctive feature being an uncharacteristically large VirB7 lipoprotein subunit [17] whose C-terminal N0 domain decorates the periphery of the outer membrane layer of the core complex [18].

VirD4 and its orthologs play a key role by recognizing substrates on the cytoplasmic face of the inner membrane and directing them for secretion through the T4SS channel [9, 19-21]. A yeast two-hybrid screen using *X. citri* VirD4 as bait identified several prey proteins (initially termed XVIPs for *<u>X</u> anthomonas* <u>V</u>irD4 interacting <u>p</u>roteins) containing a conserved C-terminal domain named XVIPCD (XVIP <u>c</u>onserved <u>d</u>omain) [22]. These proteins were later shown to be toxic antibacterial effectors secreted via the *X. citri* T4SS into target cells, often carrying N-terminal domains with enzymatic activities predicted to target structures in the cell envelope, including peptidoglycan-targeting glycohydrolases and proteases, phospholipases, as well as nucleases [6]. Furthermore, each T4SS effector is co-expressed with a cognate immunity protein, which functions to prevent self-intoxication [6], a feature also observed for toxin-antitoxin pairs associated with T6SSs [23]. Bioinformatic analysis identified potential XVIPCD-containing proteins in many other bacterial species of the Xanthomonadaceae family, including *Stenotrophomonas* spp., *Lysobacter* spp., *Luteimonas* spp., and species of the closely related Rodanobacteraceae family, including *Luteibacter* spp. and *Dyella* spp. Therefore, these effectors and their cognate immunity proteins were generally designated X-Tfes and X-Tfis (Xanthomonadaceae T4SS effectors and immunity proteins, respectively) [6].

*Stenotrophomonas maltophilia* is an emerging multi-drug-resistant global opportunistic pathogen. *S. maltophilia* strains are frequently isolated from water, soil and in association with plants [24], but in the last decades an increased number of hospital-acquired infections, particularly of immunocompromised patients, has called attention to this opportunistic pathogen [25, 26]. Infections associated with virulent strains of *S. maltophilia* are very diverse, ranging from respiratory and urinary tract infections to bacteremia and infections associated with intravenous cannulas and prosthetic devices [25]. The ability of *Stenotrophomonas* spp. to form biofilms on different biotic and abiotic surfaces [27, 28] and its capacity to secrete several hydrolytic enzymes (proteases, lipases, esterases) that promote cytotoxicity contribute to pathogenesis [29, 30]. In addition, *S. maltophilia* is naturally competent to acquire foreign DNA, which probably contributes to the multi-drug-resistant phenotype of several strains [24, 31].

*S. maltophilia* strain K279a contains a cluster of genes on its chromosome encoding for a T4SS homologous to the T4SS of the plant pathogen *Xanthomonas citri* involved in interbacterial antagonism [6], and their cytoplasmic ATPases VirD4 share 79% amino acid identity (Figure 1A). In this study, we show that *S. maltophilia* K279a is proficient in inducing *Escherichia coli* death in a T4SS-dependent manner. Interestingly, *S. maltophilia* and *X. citri* can duel using their T4SSs and kill each other. We identified eleven putative new effectors (X-Tfes) encoded by the *S. maltophilia* T4SS genome. Further characterization of one of the effectors (Smlt3024) demonstrated that it is secreted by *Stenotrophomonas* K279a in a T4SS-dependent manner upon contact with *E. coli*. Additionally, the heterologous expression of Smlt3024 in the periplasm of *E. coli* had a deleterious effect on its growth and altered cell size, which could be neutralized by co-expression of the respective cognate immunity protein, Smlt3025. This work expands our current knowledge about the subtypes of T4SSs that are involved in interbacterial competition and about the molecular mechanisms contributing to *S. maltophilia* establishment in hospital and environmental settings that may contribute to its behavior as an opportunistic pathogen.

**Fig 1.**
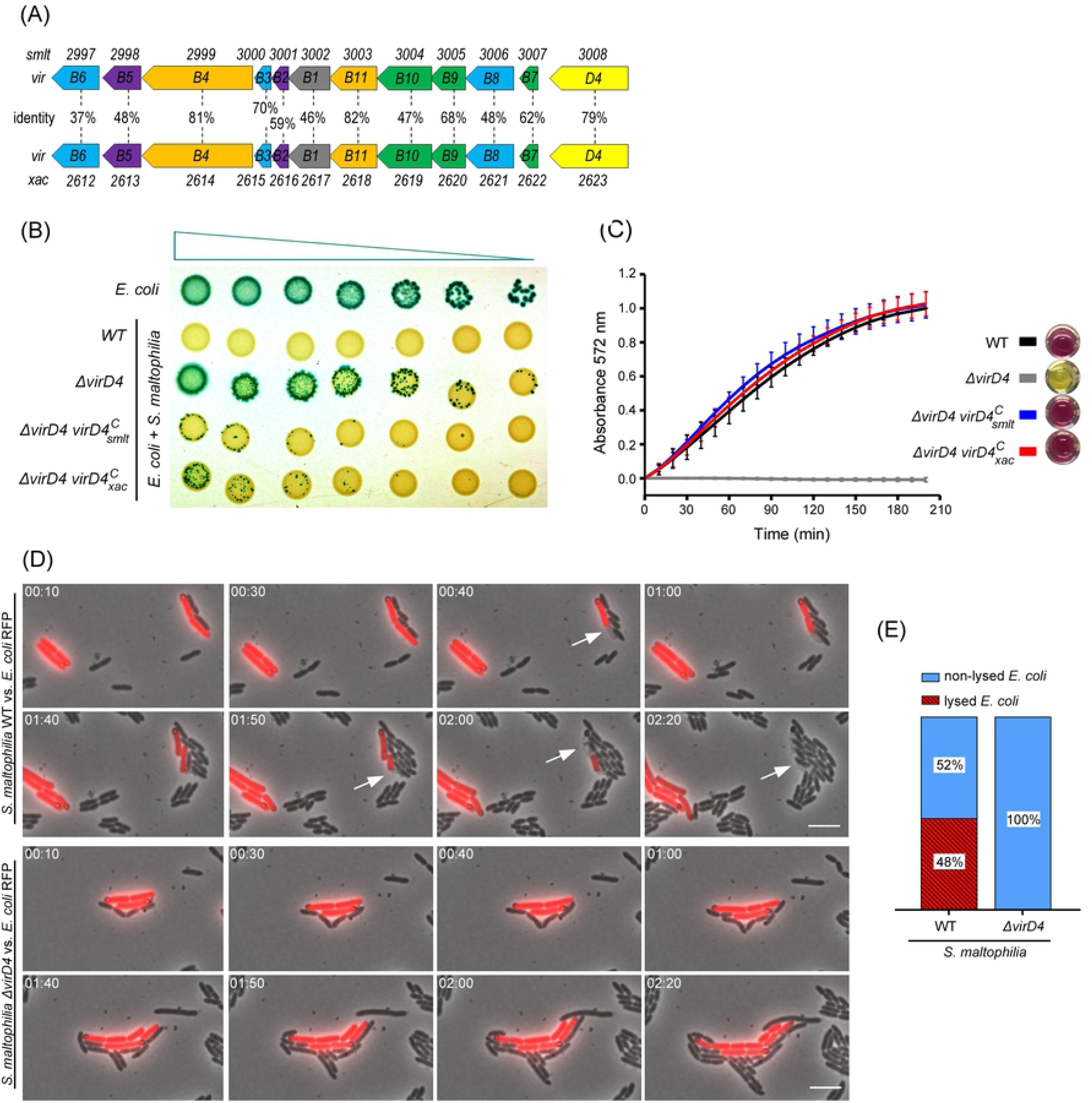
*S. maltophilia* uses the VirB/T4SS to induce *E. coli* cell death. (A) Schematic representation of the organization of the chromosomal *virB1-11* and *virD4* genes encoding the VirB/T4SSs of *S. maltophilia* K279a and *X. citri* 306. The amino acid sequence identities (%) between homologs are shown. (B) Bacterial competition assay using *S. maltophilia* strains (wild-type, Δ*virD4* and complemented strains Δ*virD4 virD4*^*C*^_*smlt*_ and Δ*virD4 virD4*^*C*^_*xac*_ and *E. coli* (naturally expressing β-galactosidase). A serial dilution of *E. coli* (1:4) was mixed with constant amounts of *S. maltophilia*, spotted onto LB-agar containing IPTG and X-gal and incubated for 24 h at 30°C. Representative image of three independent experiments. (C) Quantification of *E. coli* target cell lysis using the cell-impermeable compound CPRG. The same bacterial strains described in (B) were used. Graph represents the means and standard deviation (SD) of three independent experiments performed in triplicate. (D) Representative images of time-lapse microscopy showing wild-type *S. maltophilia* interacting with *E. coli*-*RFP* (upper panel) at the single cell level. Images were acquired every 10 min. Dead/lysed *E. coli* cells are indicated by white arrows. Interaction between *S. maltophilia* Δ*virD4* and *E. coli*-*RFP* strains (lower panel) did not induce target cell lysis. Timestamps in hours:minutes. Scale bar 5 μm. (E) Percentage of *E. coli* cells that died/lysed after cell-to-cell contact with *Stenotrophomonas* strains over a 100 min time-frame.

## Results

### *Stenotrophomonas maltophilia* VirB/T4SS induces target bacteria cell death

The genome of *S. maltophilia* K279a [32] harbours two clusters of genes encoding distinct T4SSs: *smlt1283-smlt1293* (annotated as *trb*) and *smlt2997-smlt3008* (annotated as *virB*) [33]. Comparative sequence analysis showed that the *S. maltophilia virB1-11 and virD4* genes are most closely related with their counterparts in *X. citri* involved in bacterial killing (37% - 82% identity at the amino acid level), with the three ATPases that energize the system presenting the greatest levels of identity: VirB4 (81%), VirB11 (82%) and VirD4 (79%) (Fig. 1A). Phylogenetic analysis based on the amino acid sequences of *S. maltophilia* VirD4/Smlt3008 grouped the *S. maltophilia* VirB/T4SS together with the *X. citri* VirB/T4SS involved in bacterial killing, while *Stenotrophomonas* Trb/T4SS belongs to another group of T4SSs (S1 Fig). The second T4SS from *X. citri* (encoded by plasmid pXAC64), which was proposed to be involved in conjugation due to neighbouring relaxosome genes and *oriT* site, is located in another branch in the phylogenetic tree, distinctly from the above two systems (S1 Fig) [22].

To investigate the involvement of the *S. maltophilia* VirB/T4SS in bacterial antagonism, we created a mutant strain lacking the ATPase coupling protein VirD4 (Δ*virD4*) and analysed its ability to restrict growth of other species such as *E. coli*. Different dilutions of an *E. coli* culture were mixed with a fixed number of *S. maltophilia* cells and the co-cultures were spotted onto LB-agar plates containing the chromogenic substrate X-gal and incubated for 24 h at 30°C (Fig 1B). As only *E. coli* cells naturally express β-galactosidase, they turn blue while *S. maltophilia* cells are yellow. Growth of *E. coli* was inhibited by *S. maltophilia* wild-type, but not by the Δ*virD4* strain (Fig 1B). The phenotype of *S. maltophilia* Δ*virD4* could be restored by complementing the strain with a plasmid encoding VirD4 (*smlt3008*) under the control of the P_BAD_ promoter (Δ*virD4 virD4*^*C*^_*smlt*_) (Fig 1B). This plasmid promotes low expression levels under non-inducing conditions (no L-arabinose) in *Stenotrophomonas* and is usually sufficient for complementation. Interestingly, transformation of *S. maltophilia* Δ*virD4* strain with a plasmid encoding VirD4 from *X. citri* (*xac2623*) (Δ*virD4 virD4*^*C*^_*xac*_) also restored the phenotype, indicating that these proteins are functionally redundant (Fig 1B).

To analyse bacterial antagonism at shorter time-points, we measured *E. coli* cell lysis after mixing with different *S. maltophilia* strains (wild-type, Δ*virD4*, Δ*virD4 virD4*^*C*^_*smlt*_ and Δ*virD4 virD4*^*C*^_*xac*_). The cultures were mixed and immediately spotted onto 96 well plates containing LB-agar with CPRG, which is a cell-impermeable chromogenic substrate hydrolysed by β-galactosidase released from lysed *E. coli*, thus producing chlorophenol red with an absorbance maximum at 572 nm. Fig 1C shows that *S. maltophilia* wild-type and complemented strains (Δ*virD4 virD4*^*C*^_*smlt*_ and Δ*virD4 virD4*^*C*^_*xac*_) induce lysis of *E. coli* shortly after co-incubation (around 10 min). Based on the slope of the curves, the data suggests that all three *S. maltophilia* strains induce *E. coli* cell lysis with very similar efficiencies (Fig 1C).

Live-cell imaging of *S. maltophilia* wild-type co-incubated with *E. coli-RFP* expressing red fluorescent protein (RFP) shows that *Stenotrophomonas* induces *E. coli* cell lysis in a contact-dependent manner (Fig 1D and Movie S1). No cell lysis was detected when *E. coli* was co-incubated with *S. maltophilia* Δ*virD4* (Fig 1D and Movie S2). Quantification of *E. coli* cell lysis over a time-frame of 100 min shows that around 50% of *E. coli* cells in contact with *Stenotrophomonas* wild-type were observed to lyse during this period, while no cell lysis was detected when *E. coli* was mixed with *S. maltophilia* Δ*virD4* (Fig 1E). It is important to note that this quantification does not measure (most likely sub-estimates) the efficiency of killing as some *E. coli* cells may be intoxicated without cellular lysis and the time of target-cell lysis may vary after the initial physical contact.

As *X. citri* is the only other bacterial species described to date experimentally shown to use a T4SS for interbacterial killing, we decided to analyse whether *S. maltophilia* and *X. citri* could use their T4SS to compete with and kill each other. First, we co-incubated *S. maltophilia* (either wild-type or Δ*virD4)* with a *X. citri* T4SS mutant strain lacking all the chromosomal *virB* genes and expressing green fluorescent protein (GFP) under the control of *virB7* endogenous promoter (Δ*virB-GFP*) (Cenens et al., manuscript in preparation) and confirmed that *S. maltophilia* can induce lysis of *X. citri* Δ*virB*-*GFP* in a T4SS-dependent manner (Figs 2A and 2C; Movie S3 and S4). Next, we co-incubated *X. citri*-*GFP* (functional T4SS) with *S. maltophilia* wild-type or Δ*virD4* strains. Besides showing that *X. citri* can induce lysis of *S. maltophilia* Δ*virD4* (Movie S5), we observed that when both wild-type species are mixed, they duel and kill each other in a T4SS-dependent manner (Figs 2B and 2D; Movie S6). *S. maltophilia* seems to be slightly more effective in killing *X. citri* via its T4SS, which could be due to the efficiency of the T4SS and/or the shorter doubling-time of *S. maltophilia* compared to *X. citri*.

**Fig 2.**
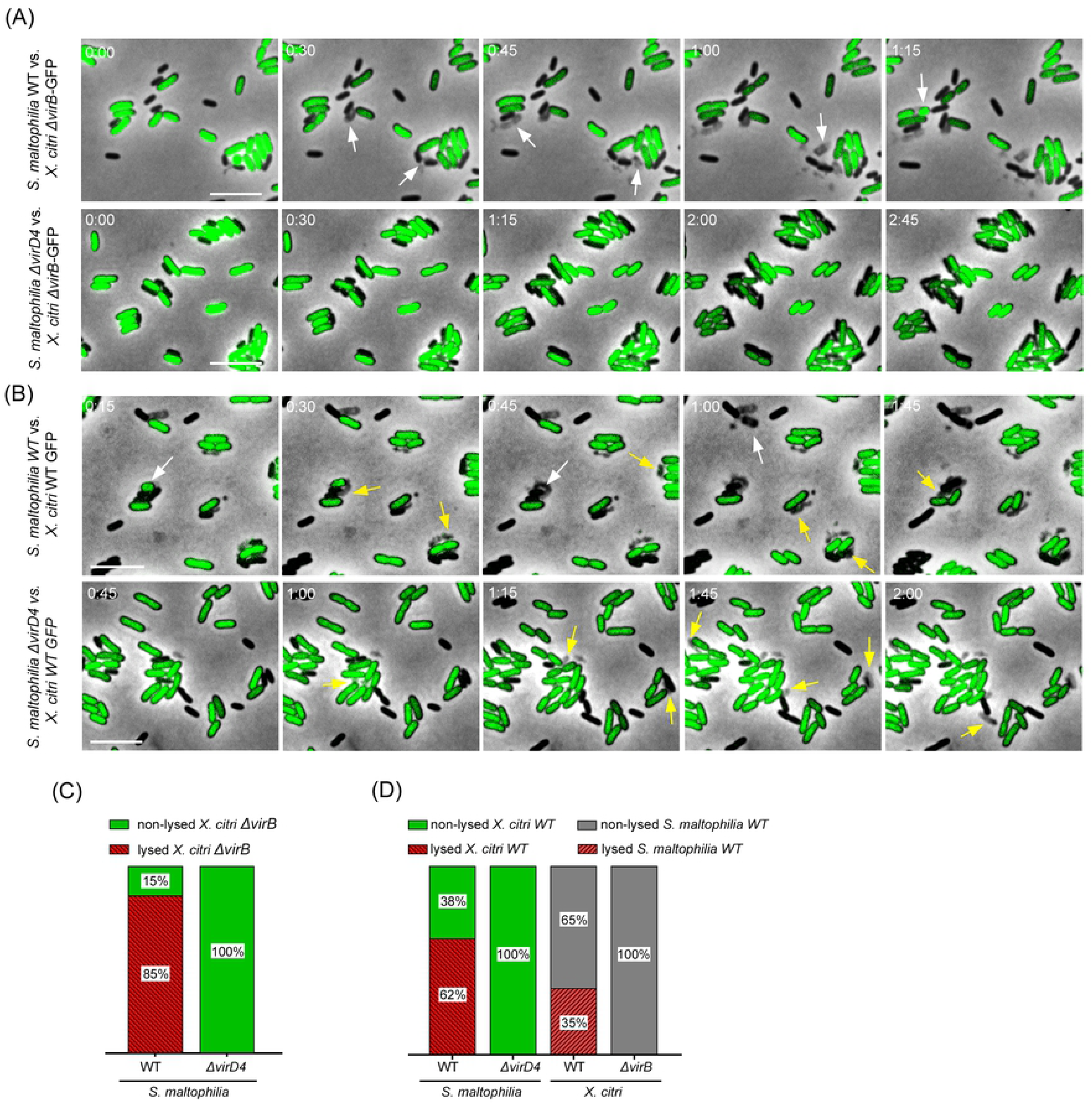
*S. maltophilia* and *X. citri* use their T4SSs for reciprocal interbacterial competition. (A) Representative images of time-lapse microscopy showing *S. maltophilia* wild-type and Δ*virD4* strains interacting with the T4SS-deficient *X. citri* Δ*virB*-*GFP* strain at the single cell level. Dead/lysed *X. citri* Δ*virB*-*GFP* cells are indicated by white arrows. (B) *S. maltophilia* wild-type and Δ*virD4* strains interacting with the functional T4SS *X. citri*-*GFP* strain at the single cell level. Dead/lysed *X. citri* cells are indicated by white arrows and dead/lysed *S. maltophilia* cells are indicated by yellow arrows. Scale bar 5 μm. Images were acquired every 15 min. (C) Percentage (%) of T4SS-deficient *X. citri* cells that lysed after cell-to-cell contact with *S. maltophilia* cells. (D) Percentage (%) of *X. citri* cells that lysed after cell-to-cell contact with *S. maltophilia* cells (left side) and % of *S. maltophilia* cells that lysed after cell-to-cell contact with *X. citri* cells (right side). Cells were counted per interaction over a 300 min time-frame.

### Identification of eleven putative effectors secreted via the *S. maltophilia* VirB/T4SS

After confirming that the *S. maltophilia* VirB/T4SS is functional and induces target cell death, we decided to search for effector proteins translocated by this system. As the VirD4 coupling protein of *X. citri* complements the Δ*virD4* strain of *S. maltophilia* (Figs 1B and 1C), we hypothesized that potential substrates secreted via the T4SS of *S. maltophilia* could be identified by applying a bioinformatic approach using the C-terminal domains of *X. citri* X-Tfes (XVIPCDs) to search the genome of *S. maltophilia* K279a. Using this approach, we identified eleven *S. maltophilia* proteins as potential T4SS substrates (Fig 3A, S1 Table). All these putative X-Tfes were identified using searches with the XVIPCDs from at least seven different *X. citri* X-Tfes (S1 Table). Amino acid sequence alignment of C-terminal XVIPCDs from *Stenotrophomonas* X-Tfes revealed a series of conserved amino acid motifs that are also present in *X. citri* X-Tfes [22] (Fig 3B).

**Fig 3.**
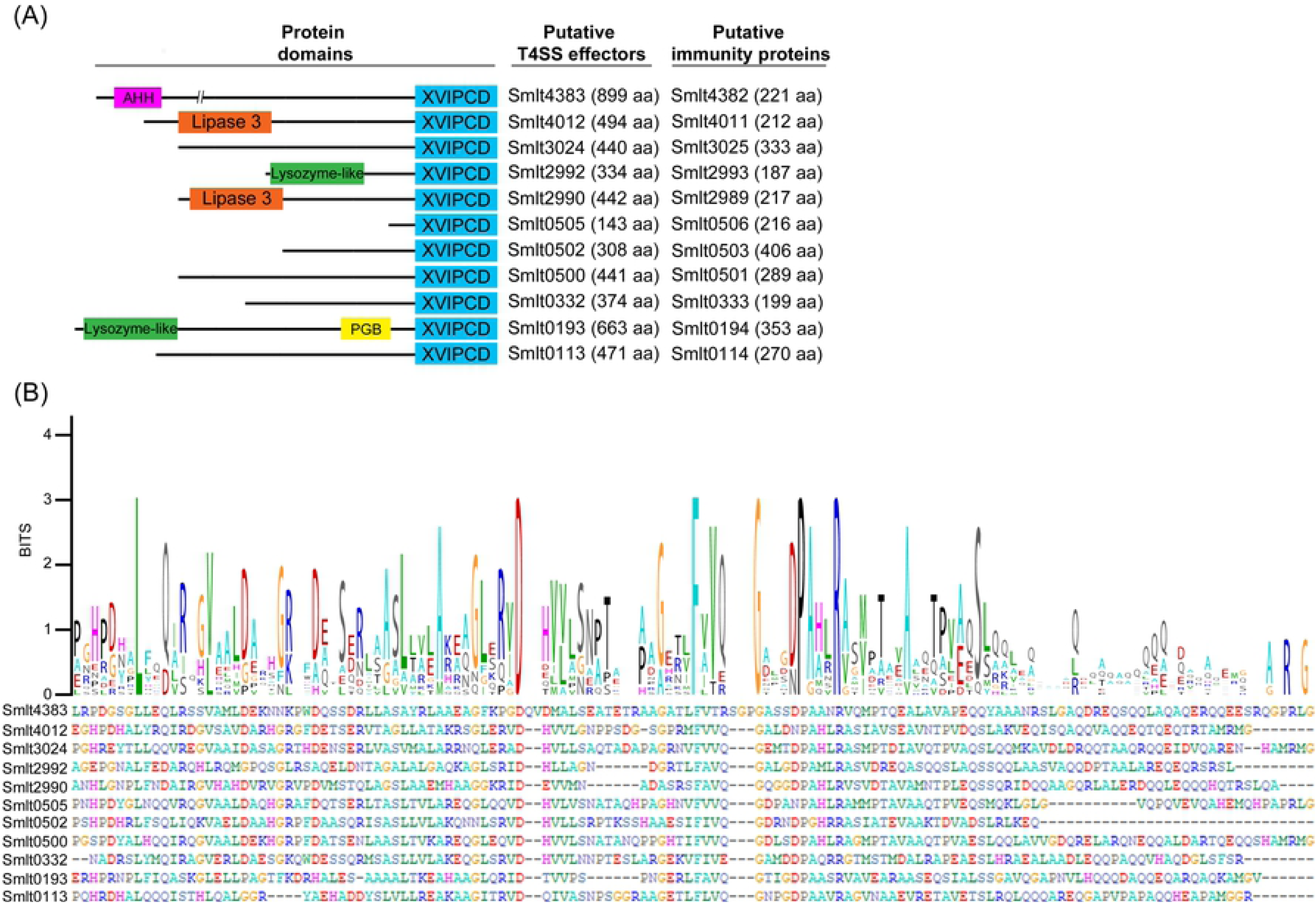
New putative type 4 effectors (Tfes) and type 4 immunity proteins (Tfis) of *S. maltophilia* T4SS. (A) Schematic representation of size and domain architectures of new *S. maltophilia* T4SS substrates identified via BLASTp search using XVIPCD (*Xanthomonas* VirD4-interacting protein conserved domain) of *X. citri* T4SS effectors. Gene entries are shown for both effectors and immunity proteins. AHH: putative nuclease domain; PGB: peptidoglycan-binding domain. (B) Alignment of the XVIPCDs of the identified *S. maltophilia* effectors using Clustal Omega [62] and consensus sequence logo generated by WebLogo [63] showing several highly conserved amino acids that match conserved residues of the *X. citri* XVIPCDs [22].

All identified *S. maltophilia* effectors are organized in small operons together with an upstream gene encoding a conserved hypothetical protein, reminiscent of the organization of effectors with their immunity proteins [6, 34]. Five of the identified *S. maltophilia* T4SS substrates harbour domains already described in other bacterial toxins such as lipases, nucleases, lysozyme-like hydrolases and proteins with peptidoglycan binding domains (Fig 3A). Three of these effectors (*smlt2990, smlt2992* and *smlt3024*) are encoded by genes very close to the *S. maltophilia virB* locus (genes *smlt2997* to *smlt3008*), further illustrating the link of these effectors with the T4SS. It is interesting to note that six of the identified putative *Stenotrophomonas* T4SS effectors do not display any known protein domain that could indicate the mechanism mediating the antibacterial toxicity (*smlt0113, smlt0332, smlt0500, smlt0502, smlt0505, smlt3024*) (Fig 3). To validate our findings and obtain further insight regarding the function of the effectors with domains of unknown function, we selected the products of the *smlt3024* gene and its upstream, putatively co-transcribed, partner (*smlt3025)* for further characterization.

### Smlt3024 is a periplasmic-acting toxin that impairs cell growth and is inhibited by periplasmic Smlt3025

BLASTp searches for Smlt3024 homologues (excluding the C-terminal XVIPCD) retrieved sequences from *Stenotrophomonas* spp., *Xanthomonas* spp., and *Lysobacter* spp. annotated either as hemolysin/hemolysin-related or as hypothetical proteins (S1B Table). However, none of these proteins contain a domain similar to any annotated domain in the Pfam database [35]. In its genomic context, *smlt3024* seems to be organized in an operon downstream to two genes encoding for its putative cognate immunity protein (*smlt3025*) and another small protein containing a helix-turn-helix (HTH) domain annotated as a putative transcriptional regulator (*smlt3026*) (Fig 4A).

**Fig 4.**
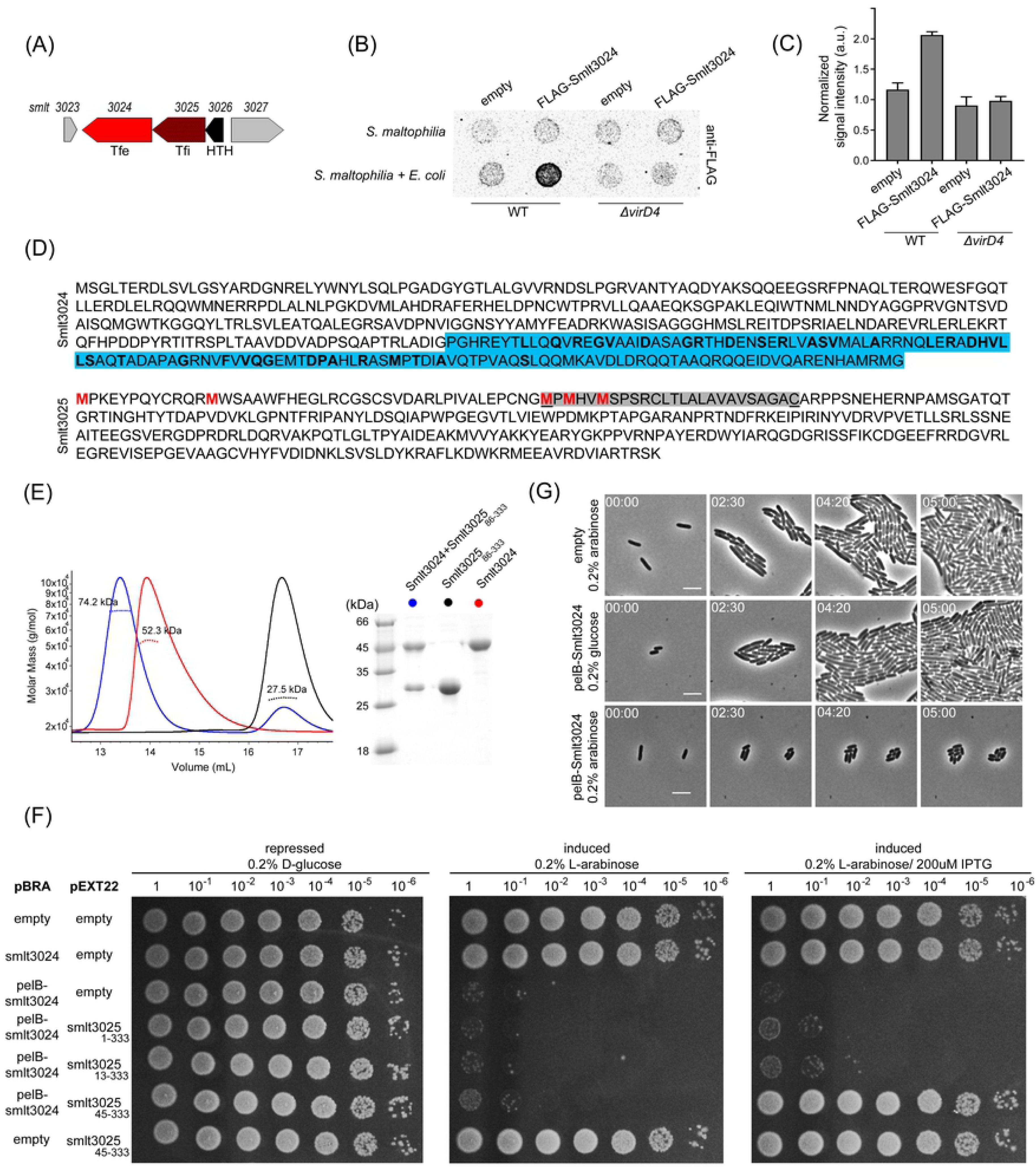
*Smlt3024* and *smlt3025* encode a Tfe-Tfi pair of *S. maltophilia* VirB/T4SS. (A) Schematic representation of *smlt3024* and *smlt3025* genomic organization. (B) Secretion assay showing T4SS-dependent and *E. coli* contact-dependent secretion of FLAG-Smlt3024. Representative image of three independent experiments. (C) Densitometry of quantitative dot blot analysis signals shown in (B). Signal intensity detected for *S. maltophilia* mixed with *E. coli* were normalized by the background signal detected for *S. maltophilia* alone. (D) Amino acid sequence of X-Tfe/X-Tfi pair Smlt3024 and Smlt3025 as annotated in *S. maltophilia* str. *K279a* genome (GenBank AM743169). *Top panel*: The amino acid sequence of Smlt3024. Coloured in blue is the XVIPCD with the most conserved amino acids in bold. Methionine (M) residues at positions 1, 13, 45, 47 and 50 are shown in red. Note that experimental evidence presented in this study suggests that the correct start codon is located at Met45, Met47 or Met50. The predicted periplasmic localization peptide of Smlt3025 beginning at Met45 is shaded in grey with cleavage predicted at the underlined cysteine. (E) Left panel: SEC-MALS analysis showing the formation of a stable complex between Smlt3024 and Smlt3025_86-333_. The continuous line corresponds to the normalized differential refractive index, and the spotted lines indicate the calculated molecular mass. *Right panel*: SDS-PAGE showing the apparent molecular mass of proteins eluted from different SEC peaks. (F) Serial dilution (10-fold) of *E. coli* strains containing pBRA and pEXT22 constructs as indicated, spotted on LB-agar plates containing either 0.2% D-glucose, 0.2% L-arabinose, 200 μM IPTG or both, showing growth inhibition of periplasmic Smlt3024 and the three versions of Smlt3025 starting in three different start codons (Smlt3025, Smlt3025_13-333_ and Smlt3025_45-333_). (G) Time-lapse imaging of single cells expressing pBRA-*pelB*-*smlt3024* show reduced growth rates and smaller cell-sizes (L-arabinose) compared to the non-induced (D-glucose) and empty plasmid controls. Images were acquired every 10 min. Timestamps in hours:minutes. Scale bar 5 μm.

To determine whether Smlt3024 is indeed an effector secreted via the *S. maltophilia* T4SS, we cloned an N-terminal FLAG-tagged version of *smlt3024* (FLAG-Smlt3024) into the pBRA plasmid under the control of the P_BAD_ promoter and used it to transform both *S. maltophilia* wild-type and Δ*virD4* strains. These strains were co-incubated with *E. coli* and spotted onto nitrocellulose membranes placed over LB-agar plates containing 0.1% L-arabinose and incubated for 6 h at 30°C. The membranes were later processed for immunodetection with an anti-FLAG antibody. Results show an increase in signal intensity for FLAG-Smlt3024 when *S. maltophilia* was co-incubated with *E. coli* (Figs 4B and 4C), while no increase was detected when *S. maltophilia* Δ*virD4* was co-incubated with *E. coli* (Figs 4B and 4C). In addition, no increase in signal intensity could be detected when *S. maltophilia* FLAG-Smlt3024 was incubated without target *E. coli* cells (Fig 4B). SDS-PAGE of total protein extracts followed by western blot with anti-FLAG antibody showed that both *S. maltophilia* wild-type and Δ*virD4* strains were expressing similar levels of FLAG-Smlt3024 (S2 Fig). These results show that secretion of Smlt3024 is dependent on contact with a target cell and on a functional T4SS.

To assess whether Smlt3025 is the cognate immunity protein of Smlt3024, we analysed whether these proteins could interact by expressing and purifying full-length Smlt3024 and a soluble version of Smlt3025 (amino acid residues 86-333), lacking its N-terminal signal peptide. Complex formation was analysed using size exclusion chromatography coupled to multiple-angle light scattering (SEC-MALS) (Fig 4E). The MALS analysis calculated average masses for Smlt3024 and Smlt3025_86-333_ of 52.3 kDa and 27.5 kDa, respectively, which are very close to the theoretical values of their monomer molecular masses of 49 kDa and 28 kDa, respectively (Fig. 4E). When a mixture of these proteins was analysed by SEC-MALS followed by SDS-PAGE, a new peak appeared containing both Smlt3024 and Smlt3025_86-333_ with an average molecular mass calculated by MALS of 74.2 kDa, suggesting that a stable 1:1 complex (theoretical mass of 77 kDa) was formed between Smlt3024 and Smlt3025_86-333_ (Fig 4E).

If Smlt3024 is indeed a toxic effector secreted by the *S. maltophilia* T4SS, then we would expect that its expression in the appropriate compartment within *E. coli* would create an impairment of bacterial growth. To evaluate the toxicity of Smlt3024 upon expression in *E. coli* and to establish in which cellular compartment Smlt3024 exerts its effect, we cloned the full-length protein into pBRA placing it under control of the P_BAD_ promoter (inducible by L-arabinose and repressed by D-glucose) both with and without an N-terminal *pelB* periplasmic localization signal sequence. We also cloned the sequence of the Smlt3025 immunity protein into the pEXT22 vector placing it under the control of P_TAC_ promoter, which can be induced by IPTG. We noted that the annotated sequence for Smlt3025 has a non-canonical GTG start codon with 4 possible in frame ATG start codons at positions 13, 45, 47 and 50 and that initiation at positions 45, 47 or 50 would produce proteins with an N-terminal signal sequence for periplasmic localization (Fig. 4D) [36]. Therefore, three versions of Smlt3025, beginning at positions 1, 13 and 45 were cloned into pEXT22, leading to the production of Smlt3025_1-333_, Smlt3025_13-333_ and Smlt3025_45-333_. *E. coli* strains carrying the different combinations of pBRA-Smlt3024 and each one of the pEXT22-Smlt3025 plasmids were serial diluted and incubated on LB-agar plates containing either D-glucose, L-arabinose or L-arabinose plus IPTG (D-glucose inhibits and L-arabinose induces expression of Smlt3024; IPTG induces expression of Smlt3025). Results showed that Smlt3024 is toxic only when directed to the periplasm of *E. coli* cells (pBRA-*pelB-smlt3024*) but not in the cytoplasm (pBRA-*smlt3024*), and that only Smlt3025_45-333_ which is directed to the periplasm, could neutralize Smlt3024 toxicity (Fig. 4F).

To gather further insight on the mechanism by which Smlt3024 could induce toxicity, we decided to perform time-lapse microscopy to evaluate growth and morphology of individual *E. coli* cells carrying the empty pBRA or pBRA-*pelB-smlt3024* plasmids. *E. coli* carrying the empty plasmid incubated on LB-agar with 0.2% L-arabinose (Fig 4G and Movie S7) as well as the repressed pBRA-*pelB-smlt3024* (0.2% D-glucose) grew normally (Fig 4G and Movie S8). Upon induction with L-arabinose, cells carrying pBRA-*pelB-smlt3024* quickly experienced a strong reduction in growth rate and single cells were smaller (average length of 2.1 ± 0.7 μm after 300 min) compared to the controls incubated in glucose (average length of 3.6 ± 1.2 μm after 300 min) (Fig 4G and Movie S9). Despite the severe delay in doubling time, *E. coli* cells expressing PelB-Smlt3024 seem to remain viable and continue growing and dividing for up to 8 h (Movie S9).

## Discussion

Competition between microorganisms for nutrients and space often determines which species will thrive and dominate or be eradicated from a specific habitat. The recently identified T4SS involved in bacterial killing has revealed another weapon in the bacterial warfare arsenal [6]. In this manuscript, we show that the T4SS of *S. maltophilia* is involved in interbacterial competition, allowing it to induce lysis of *E. coli* and *X. citri* and possibly many other Gram-negative species. The *S. maltophilia* VirB/T4SS is the second example described to date of a T4SS subtype involved in bacterial killing and it is the first example of a human opportunistic pathogen shown to mediate bacterial killing via a T4SS.

*S. maltophilia* is often found as a member of microbial communities in water, soil and in association with plants. Some *Stenotrophomonas* species like *S. maltophilia* and *S. rhizophila* can participate in beneficial interactions with plants, but no species were reported to be phytopathogenic, which distinguishes *Stenotrophomonas* from the phylogenetically related genera *Xanthomonas* and *Xylella* [24]. More importantly, an increasing number of hospital-acquired *S. maltophilia* infections over the last decades has led to the classification of this bacterium as an emerging opportunistic pathogen [25, 26]. Key to the opportunistic behaviour of *S. maltophilia* strains are their ability to form biofilms and their resistance to multiple antibiotics. In this context, the antibacterial property of its T4SS probably provides a competitive advantage to *S. maltophilia* in polymicrobial communities, contributing to increased fitness.

The most worrying aspect of pathogenic *S. maltophilia* strains is their multi-drug resistance phenotype [37]. As *S. maltophilia* is naturally competent to acquire foreign DNA (Berg et al., 1999, Ryan et al., 2009), it would be interesting to analyse the contribution of the T4SS described here in promoting *Stenotrophomonas* transformation and the acquisition of antibiotic resistance genes. Such a mechanism has already been reported in *Vibrio cholerae*, which uses a T6SS as a predatory killing device to induce target cell lysis concomitantly with target-cell DNA uptake to promote bacterial transformation [38].

The T6SS has been shown to play important roles during colonization and infection by enteric pathogenic bacteria such as *Salmonella* and *Shigella* [39, 40]. The contribution of *S. maltophilia* T4SS to colonization and maintenance within mammalian hosts is still unknown. As *S. maltophilia* is frequently associated with cystic fibrosis patients [41, 42], it would be interesting to analyse the ability of mutant strains lacking a functional T4SS to compete with oral and nasal microbiota during infection of susceptible model organisms [43, 44].

The new effector/immunity protein (X-Tfe/X-Tfi) pairs identified in this study expand our knowledge on the different effector proteins promoting target-cell toxicity mediated by T4SSs. Most of the characterized T4SS and T6SS antibacterial toxins are enzymes that degrade structural cellular components such as peptidoglycan and phospholipids, thus promoting target cell lysis [45]. Recent studies have identified effectors that change cell metabolism promoting altered cell growth rather than lysis [46, 47]. Promoting target cell stasis is in most cases sufficient to provide the attacker with a competitive advantage, allowing it to outnumber the target species and establish itself in the environment. In natural settings, many species are likely to have acquired resistance mechanisms by means of immunity proteins against specific secreted effectors and may be sensitive to only one or a few effectors within the secreted cocktail. Additionally, bacterial effectors work synergistically and display conditional efficiency depending on the environment [48]. An example of the diversity of effector-immunity pairs carried by different organisms is clearly illustrated here by the duelling observed between *S. maltophilia* and *X. citri*, which can kill one another in a T4SS-dependent manner, indicating that each species lacks at least one immunity protein against the rival’s set of T4SS effectors.

Among the eleven new *S. maltophilia* T4SS effectors identified by bioinformatic searches, six of them have N-terminal domains that are not significantly similar in sequence to any other annotated protein domain family. These X-Tfes are particularly interesting since they could possibly promote toxic effects by different mechanisms from those described for the majority of antibacterial type 6- and type 4 effectors that target the cell wall, membranes or nucleic acids [6, 46, 49-51]. Smlt3024 induced a severe reduction in growth speed and a decrease in cell size in rich media. However, despite the delay, cells continue to grow for over 8 h (Movie S9), indicating that Smlt3024 expression does not lead to cell death in laboratory conditions. As slower growth speed and decreased cell-sizes are reminiscent of cells growing in nutrient deprived media [52], we speculate that Smlt3024 could somehow induce a starvation response either by mimicking a systemic response, blocking an important metabolic pathway or blocking a key nutrient importer.

This work contributes with knowledge on the virulence mechanisms used by *S. maltophilia* allowing it to survive in polymicrobial communities and maintain environmental reservoirs. The work also expands our current knowledge about the subtypes of T4SSs involved in interbacterial killing for which the diversity and mechanism of toxicity from secreted substrates, and distribution among bacterial species we are only beginning to understand.

## Materials and Methods

### Bacterial strains and culture conditions

*S. maltophilia* K279a [32] and *X. citri pv. citri* 306 [53] were grown in 2x YT media (16 g/L tryptone, 10 g/L yeast extract, 5 g/L NaCl). *E. coli* strain K-12 subsp. MG1655 [54] was used in competition assays because of its endogenous expression of β-galactosidase. *E. coli* DH5α and *E. coli* HST08 were used for cloning purposes and *E. coli* S17 was used for conjugation with *S. maltophilia*. The *X. citri* T4SS mutant strain lacks all chromosomal *virB* genes and has the *msfGFP* gene under the control of *virB7* endogenous promoter (Δ*virB*-*GFP*) while the *X. citri-GFP* strain has a functional T4SS and expresses GFP as transcriptional fusion under the control of *virB7* promoter (Cenens et al., in preparation). For time-lapse imaging of *S. maltophilia* and *X. citri* strains, AB defined media was used (0.2% (NH_4_)_2_SO_4_, 0.6% Na_2_HPO_4_, 0.3% KH_2_PO_4_, 0.3% NaCl, 0.1 mM CaCl_2_, 1 mM MgCl_2_, 3 μM FeCl_3_) supplemented with 0.2% sucrose, 0.2% casamino acids, 10 μg/mL thiamine and 25 μg/mL uracil. Cultures of *E. coli* and *S. maltophilia* were grown at 37°C with agitation (200 rpm) and *X. citri* cultures were grown at 28°C with agitation (200 rpm). Antibiotics were used at the following concentrations to select *S. maltophilia* strains: tetracycline 40 μg/mL and streptomycin 150 μg/mL. For selection of *E. coli* strains, kanamycin 50 μg/ml and spectinomycin 100 μg/ml were used when appropriate. For induction from P_BAD_ promoter, 0.2% L-arabinose was added. For P_TAC_ induction, 200 μM IPTG was used. Expression from both promoters was repressed using 0.2% D-glucose.

### Cloning and mutagenesis

All primers and plasmids used for cloning are listed in S2 Table. To produce in-frame deletions of *virD4* (*smlt3008*), we used a two-step integration/excision exchange process and the pEX18Tc vector [55]. Fragments of ∼1000-bp homologous to the upstream and downstream regions of *smlt3008* were amplified by PCR and cloned into pEX18Tc using standard restriction digestion and ligation. The pEX18Tc-Δ*virD4* was transformed in *E. coli* S17 donor cells by electroporation and transferred to *S. maltophilia* recipients via conjugation following the protocol described by Welker et al. [56]. Tetracycline-resistant colonies were first selected. Colonies were then grown in 2x YT without antibiotic and plated on 2x YT agar containing 10% sucrose without antibiotic. Mutant clones were confirmed by PCR. To complement the Δ*virD4* strain, the gene encoding full-length *smlt3008* was PCR amplified from genomic DNA and cloned into the pBRA vector, which is a pBAD24-derived vector that promotes low constitutive expression in *Stenotrophomonas*/*Xanthomonas* under non-inducing conditions (M. Marroquin, unpublished). The pBRA construct encoding full-length *X. citri virD4*/*XAC2623* was reported previously [6]. For secretion assays, the full-length sequence of *smlt3024* was cloned into pBRA vector, including a FLAG tag on its N-terminus and transformed into *S. maltophilia* wild-type and Δ*virD4*. Plasmids were transformed into *S. maltophilia* by electroporation (2.5 kV, 200 Ω, 25 μF, 0.2 cm cuvettes), followed by streptomycin selection. For cloning *smlt3024* and *smlt3025* into pSUMO – a modified version of pET28a (Novagen), adding a SUMO tag between the hexahistidine and the cloning site – we used the soluble portion of Smlt3025 (residues between 86-333) that lacks the N-terminal signal peptide and the full-length Smlt3024. To produce *smlt3024* with the *pelB* periplasmic localization sequence, PCR products were first cloned in pET22b (Novagen; containing the N-terminal *pelB* sequence) after the *pelB-smlt3024* construct was transferred to pBRA using Gibson assembly. For the immunity protein *smtl3025*, three different constructs were cloned in pEXT22 [57]: one starting at the annotated start-codon and two starting at two downstream putative start-codons (positions 17 and 45). The sequences of all constructs containing effectors in pBRA and immunity proteins in pEXT22 were confirmed by DNA sequencing to assure absence of point mutations in the cloned genes and upstream promoter sites using the Macrogen standard sequencing service (https://dna.macrogen.com/).

### Bacterial competition assays

Bacterial competition was assessed either by analysing target cell growth or target cell lysis. To analyse *E. coli* growth during co-incubation with *S. maltophilia* we used a protocol adapted from Hachani et al. [58]. Briefly, strains were subcultured 1:100 and grown to exponential phase for 2 h at 37°C (200 rpm). Cells were washed with 2x YT and the optical density measured at 600 nm (OD_600nm_) adjusted to 1. Serial dilutions (1:4) of *E. coli* culture was performed in 96 well plates. Equal volumes of *E. coli* and *S. maltophilia* cultures at OD_600nm_ 1.0 were mixed into each well. After mixing, 5 μl were spotted onto LB-agar plates containing 100 μM IPTG (isopropyl β-D-1-thiogalactopyranoside) and 40 μg/mL X-gal (5-bromo-4-chloro-3-indolyl-β-D-galactopyranoside) using multichannel pipettes. Plates were incubated for 24 h at 30°C. Analysis of target cell death was performed using CPRG (chlorophenol red-β-D-galactopyranoside) as described previously with minor modifications [18, 59]. Briefly, *S. maltophilia* and *E. coli* overnight cultures were subcultured 1:100 – the latter containing 200 μM IPTG – and grown at 37°C (200 rpm) to reach OD_600nm_ of approximately 1. Cells were washed with LB media, OD_600nm_ adjusted to 1.0 for *S. maltophilia* strains and OD_600nm_ adjusted to 8.0 for *E. coli*. The adjusted cultures were mixed 1:1 and 10 μL spotted in triplicate onto 96 well plates containing 100 μL of solid 1.5% 2x YT agar and 40 μg/mL CPRG. Plates were let dry completely, covered with adhesive seals and analysed on a SpectraMax Microplate Reader (Molecular Devices) at 572 nm every 10 min for 3.5 h. *E. coli* cultures were also spotted onto the same plate as a control for spontaneous cell death. The obtained A_572_ data was processed using RStudio (www.rstudio.com) and plotted using the ggplot2 package [60]. Background intensities obtained from the mean A_572_ values containing only *E. coli* cells were subtracted from all data series. The initial A_572_ value at time-point 0 min was subtracted from all subsequent time-points to correct for small differences in initial measurements. Finally, the curves of *S. maltophilia* Δ*virD4* and complementation strains were normalized to those obtained for the *S. maltophilia* wild-type strain.

### Time-lapse microscopy

For time-lapse imaging of bacterial killing at the single-cell level, agar slabs containing either 2x YT or supplemented AB media were created by cutting a rectangular frame out of a double-sided adhesive tape (3M™ VHB™ transparent, 24 mm wide, 1 mm thick), which was taped onto a first microscopy slide. Into the resulting tray, agar was poured and covered by a second microscopy glass slide to create a smooth surface. After solidification, the second microscopy slide was removed, exposing the agar’s surface onto which 2 μl of cell suspensions were spotted. After cell suspensions were left to dry completely, a #1.5 cover glass (Corning) was laid on top of the agar slab and closed at the sides by the second adhesive layer of the tape, leaving the cell mixtures closely and stably pressed between cover glass and the agar slab. Soon after, phase contrast images together with GFP or RFP excitation images were obtained with a Leica DMI-8+ epi-fluorescent microscope equipped with the Leica DFC365 FX camera, a HC PL APO 100x/1.4 Oil ph3 objective (Leica), a GFP excitation-emission band-pass filter cube (Ex: 470/40, DC: 495, EM: 525/50; Leica) and a Cy3/Rhodamine excitation-emission band-pass filter cube (Ex: 541/51, DC: 560, EM: 565/605; Leica). An incubation cage around the microscope kept temperatures constant at 37°C for *E. coli* and *S. maltophilia* experiments and at 28°C for experiments with *X. citri*. Several separate positions of each cell mixture were imaged every 10-15 min after auto-focusing using the LASX software package (Leica). Images were further processed with the FIJI software using the Bio-Formats plugin [61]. Time-lapse images were visually scored for cell lysis events. Small groups of cells (approximately 2 to 8 cells per colony) containing a mixture of bacterial species in close contact with each other were tagged at time-point zero and followed during 100 min (*E. coli* and *S. maltophilia* competitions) or 300 min (*X. citri* and *S. maltophilia* competitions) and cell lysis events were manually registered. Approximately 100 cells were scored for each assay.

### BLASTp searches

To identify putative effectors secreted by the *S. maltophilia* T4SS, we used the XVIPCDs of known and putative *X. citri* T4SS substrates (residues in parenthesis): *XAC4264*(140-279), *XAC3634*(189-306), *XAC3266*(735-861), *XAC2885*(271-395), *XAC2609*(315-431), *XAC1918*(477-606), *XAC1165*(1-112), *XAC0574*(317-440), *XAC0466*(488-584), *XAC0323*(16-136), *XAC0151*(120-254), *XAC0096*(506-646) [6, 22] to BLAST search the genome of *S. maltophilia* K279a (https://www.genome.jp/tools/blast/). A list of *S. maltophilia* proteins identified by each *X. citri* XVIPCD with their respective E-values is shown in S1 Table.

### Recombinant protein expression, purification and SEC-MALS analysis

Smlt3025_86-333_ (residues between 86-333) and full-length Smlt3024 cloned into pSUMO were transformed into *E. coli* BL21(DE3) and SHuffle T7 competent *E. coli* cells (New England BioLabs), respectively, and subcultured into 2x YT medium supplemented with 50 μg/mL kanamycin at 37°C until OD_600nm_ of 0.6 and then shifted to 18°C. After 30 min, protein production was induced with 0.1 mM IPTG. After overnight expression, cells were harvested by centrifugation and resuspended in 20 mM Tris-HCl (pH 8.0), 200 mM NaCl, 5 mM imidazole and lysed by 10 passages in a French Press system. The lysate soluble fraction was loaded onto a 5 ml HiTrap chelating HP column (GE Healthcare) immobilized with 100 mM cobalt chloride and equilibrated with the lysis buffer. After the removal of unbound proteins, the protein was eluted with lysis buffer supplemented with 100 mM imidazole. Ulp1 was added to the eluted protein, followed by dialysis at 4°C for 12 h for removal of imidazol. The cleaved target proteins were purified after a second passage through the HiTrap chelating HP column immobilized with cobalt, being eluted in the unbound fraction. Molecular masses of the isolated proteins and the effector-immunity complex were determined by SEC-MALS (size-exclusion chromatography coupled to multi-angle light scattering), using a Superdex 200 10/300 GL (GE Healthcare) coupled to a Wyatt MALS detector. Graphs and the average molecular masses were generated using the ASTRA software (Wyatt), assuming a refractive index increment dn/dc = 0.185 mL/g.

### Secretion assay and immunoblot

Secretion assays were performed essentially as previously described [6]. Briefly, *S. maltophilia* wild-type and Δ*virD4* strains carrying pBRA-FLAG*-smlt3024* were grown overnight with antibiotics (150 μg/mL streptomycin), subcultured on the next day (1:25 dilution) and grown for an additional 2 h at 37°C (200 rpm). *E. coli* cells were subcultured (1:100 dilution) in a similar manner. *S. maltophilia* and *E. coli* cells were washed with 2x YT, OD_600nm_ adjusted to 1.0, mixed 1:1 volume and 5-10 μL were spotted onto dry nitrocellulose membranes, which were quickly placed onto LB-agar plates containing 0.1% L-arabinose to induce the expression of FLAG-Smlt3024. Plates were incubated at 30°C for 6 h, sufficient to allow detection of secreted proteins and before spontaneous cell death, which would produce background in the dot blot. After 6 h, membranes were washed with 5% low-fat milk diluted in PBS containing 0.02% sodium azide and processed for quantitative dot blot analysis with anti-FLAG rabbit polyclonal antibody, followed by IRDye 800CW anti-rabbit IgG (LI-COR Biosciences) and scanned using an Odyssey CLx infrared imaging system (LI-COR Biosciences). To obtain good signal to noise ratios, the membranes were washed in PBS/Tween (0.05%) at least four times for 1 h each. Quantification of signal intensity was performed using FIJI software [61].

## Acknowledgments

We are grateful to Dr. Diorge Paulo Souza for critical discussions and Dr. Gabriel U. Oka for advice on the secretion assays. We thank Dr. Robert Ryan for providing the strain, Dr. Alexandre Bruni-Cardoso for fluorescence microscope access, Dr. Beny Spira, Dr. Maria Carolina Quecine Verdi and Dr. Frederico José Gueiros for plasmids, and Dr. Andre Luis Berteli Ambrosio for the SHuffle T7 competent *E. coli*.

## Author contributions

E.B.-S. and C.S.F. conceived the study. E.B.-S., W.C., B.Y.M. and C.S.F., designed experiments. E.B.-S., W.C., B.Y.M., G.D.S. and I.V.M. performed the experiments and all of the authors analysed the data. E.B.-S. and C.S.F. wrote the manuscript.

## Supporting information

**S1 Fig. Phylogenetic distribution of *S. maltophilia* K279a T4SSs.** Maximum-likelihood tree with 1000 bootstrap replicates built with amino acid sequence of VirD4 (Smlt3008) homologs using MEGA 7.0 [64]. VirB/T4SSs from *S. maltophilia* and *X. citri* [6] involved in interbacterial competition are highlighted in orange. Trb/T4SS from *S. maltophilia* is in green and the VirB/T4SS involved in conjugation [65] encoded by the pXAC64 plasmid from *X. citri* strain 306 is in blue [22].

**S2 Fig. Loading control for secretion assay.** SDS-PAGE of total protein extracts followed by western blot of *S. maltophilia* strains carrying pBRA-FLAG-*smlt3024* or empty pBRA. RnhA (Ribonuclease HI) was used as loading control.

**S1 Movie. Time-lapse microscopy showing *S. maltophilia* wild-type interacting with *E. coli-RFP*.** Dead/lysed *E. coli* cells are indicated by white arrows. Images were acquired every 10 min. Timestamps in hours:minutes. Scale bar 5 μm

**S2 Movie. Time-lapse microscopy showing *S. maltophilia* Δ*virD4* interacting with *E. coli-RFP*.** Images were acquired every 10 min. Timestamps in hours:minutes. Scale bar 5 μm

**S3 Movie. Time-lapse microscopy showing *S. maltophilia* wild-type interacting with *X. citri* Δ*virB*-*GFP***. Dead/lysed *X. citri* cells are indicated by white arrows. Images were acquired every 15 min. Timestamps in hours:minutes. Scale bar 5 μm

**S4 Movie**. Time-lapse microscopy showing *S. maltophilia* Δ*virD4* interacting with *X. citri* Δ*virB*-*GFP*. Images were acquired every 15 min. Timestamps in hours:minutes. Scale bar 5 μm

**S5 Movie. Time-lapse microscopy showing *S. maltophilia* wild-type interacting with *X. citri*-*GFP*.** Dead/lysed *X. citri* cells are indicated by white arrows and dead/lysed *S. maltophilia* cells are indicated by yellow arrows. Images were acquired every 15 min. Timestamps in hours:minutes. Scale bar 5 μm

**S6 Movie. Time-lapse microscopy showing *S. maltophilia* Δ*virD4* interacting with *X. citri*-*GFP*.** Dead/lysed *S. maltophilia* cells are indicated by yellow arrows. Images were acquired every 15 min. Timestamps in hours:minutes. Scale bar 5 μm

**S7 Movie. Time-lapse microscopy showing *E. coli* cells containing the empty pBRA plasmid grown with 0.2% L-arabinose.** Images were acquired every 10 min. Scale bar 5 μm.

**S8 Movie. Time-lapse microscopy showing *E. coli* cells containing pBRA*-pelB-smlt3024* grown with 0.2% D-glucose.** Images were acquired every 10 min. Scale bar 5 μm.

**S9 Movie. Time-lapse microscopy showing *E. coli* cells containing pBRA*-pelB-smlt3024* grown with 0.2% L-arabinose.** Images were acquired every 10 min. Scale bar 5 μm.

**S1 Table. List of putative *S. maltophilia* T4SS X-Tfe/X-Tfi pairs identified by BLASTp search using *X. citri* XVIPCDs**.

**S2 Table. List of strains, primers and plasmids used in this study**.

## References

1. Garcia-Bayona L, Comstock LE (2018). Bacterial antagonism in host-associated microbial communities. Science 361.

2. Aoki SK, Pamma R, Hernday AD, Bickham JE, Braaten BA, Low DA (2005). Contact-dependent inhibition of growth in Escherichia coli. Science 309:1245–8.

3. Aoki SK, Diner EJ, de Roodenbeke CT, Burgess BR, Poole SJ, Braaten BA, et al. (2010). A widespread family of polymorphic contact-dependent toxin delivery systems in bacteria. Nature 468:439–42.

4. Pukatzki S, Ma AT, Revel AT, Sturtevant D, Mekalanos JJ (2007). Type VI secretion system translocates a phage tail spike-like protein into target cells where it cross-links actin. Proc Natl Acad Sci U S A 104:15508–13.

5. Pukatzki S, Ma AT, Sturtevant D, Krastins B, Sarracino D, Nelson WC, et al. (2006). Identification of a conserved bacterial protein secretion system in Vibrio cholerae using the Dictyostelium host model system. Proc Natl Acad Sci U S A 103:1528–33.

6. Souza DP, Oka GU, Alvarez-Martinez CE, Bisson-Filho AW, Dunger G, Hobeika L, et al. (2015). Bacterial killing via a type IV secretion system. Nat Commun 6:6453.

7. Grohmann E, Christie PJ, Waksman G, Backert S (2018). Type IV secretion in Gram-negative and Gram-positive bacteria. Mol Microbiol 107:455–471.

8. Cascales E, Christie PJ (2003). The versatile bacterial type IV secretion systems. Nat Rev Microbiol 1:137–49.

9. Christie PJ, Vogel JP (2000). Bacterial type IV secretion: conjugation systems adapted to deliver effector molecules to host cells. Trends Microbiol 8:354–60.

10. Sexton JA, Vogel JP (2002). Type IVB secretion by intracellular pathogens. Traffic 3:178– 85.

11. Guglielmini J, Neron B, Abby SS, Garcillan-Barcia MP, de la Cruz F, Rocha EP (2014). Key components of the eight classes of type IV secretion systems involved in bacterial conjugation or protein secretion. Nucleic Acids Res 42:5715–27.

12. Guglielmini J, de la Cruz F, Rocha EP (2013). Evolution of conjugation and type IV secretion systems. Mol Biol Evol 30:315–31.

13. Pitzschke A, Hirt H (2010). New insights into an old story: Agrobacterium-induced tumour formation in plants by plant transformation. EMBO J 29:1021–32.

14. Waksman G (2019). From conjugation to T4S systems in Gram-negative bacteria: a mechanistic biology perspective. EMBO Rep doi:10.15252/embr.201847012.

15. Christie PJ (2016). The Mosaic Type IV Secretion Systems. EcoSal Plus 7.

16. Chandran Darbari V, Waksman G (2015). Structural Biology of Bacterial Type IV Secretion Systems. Annu Rev Biochem 84:603–29.

17. Souza DP, Andrade MO, Alvarez-Martinez CE, Arantes GM, Farah CS, Salinas RK (2011). A component of the Xanthomonadaceae type IV secretion system combines a VirB7 motif with a N0 domain found in outer membrane transport proteins. PLoS Pathog 7:e1002031.

18. Sgro GG, Costa TRD, Cenens W, Souza DP, Cassago A, Coutinho de Oliveira L, et al. (2018). Cryo-EM structure of the bacteria-killing type IV secretion system core complex from Xanthomonas citri. Nat Microbiol 3:1429–1440.

19. Nagai H, Roy CR (2001). The DotA protein from Legionella pneumophila is secreted by a novel process that requires the Dot/Icm transporter. EMBO J 20:5962–70.

20. Burns DL (2003). Type IV transporters of pathogenic bacteria. Curr Opin Microbiol 6:29–34.

21. Cascales E, Christie PJ (2004). Definition of a bacterial type IV secretion pathway for a DNA substrate. Science 304:1170–3.

22. Alegria MC, Souza DP, Andrade MO, Docena C, Khater L, Ramos CH, et al. (2005). Identification of new protein-protein interactions involving the products of the chromosome- and plasmid-encoded type IV secretion loci of the phytopathogen Xanthomonas axonopodis pv. citri. J Bacteriol 187:2315–25.

23. Jamet A, Nassif X (2015). New players in the toxin field: polymorphic toxin systems in bacteria. MBio 6:e00285–15.

24. Ryan RP, Monchy S, Cardinale M, Taghavi S, Crossman L, Avison MB, et al. (2009). The versatility and adaptation of bacteria from the genus Stenotrophomonas. Nat Rev Microbiol 7:514–25.

25. Brooke JS (2012). Stenotrophomonas maltophilia: an emerging global opportunistic pathogen. Clin Microbiol Rev 25:2–41.

26. Adegoke AA, Stenstrom TA, Okoh AI (2017). Stenotrophomonas maltophilia as an Emerging Ubiquitous Pathogen: Looking Beyond Contemporary Antibiotic Therapy. Front Microbiol 8:2276.

27. Pompilio A, Crocetta V, Confalone P, Nicoletti M, Petrucca A, Guarnieri S, et al. (2010). Adhesion to and biofilm formation on IB3-1 bronchial cells by Stenotrophomonas maltophilia isolates from cystic fibrosis patients. BMC Microbiol 10:102.

28. Pompilio A, Pomponio S, Crocetta V, Gherardi G, Verginelli F, Fiscarelli E, et al. (2011). Phenotypic and genotypic characterization of Stenotrophomonas maltophilia isolates from patients with cystic fibrosis: genome diversity, biofilm formation, and virulence. BMC Microbiol 11:159.

29. DuMont AL, Karaba SM, Cianciotto NP (2015). Type II Secretion-Dependent Degradative and Cytotoxic Activities Mediated by Stenotrophomonas maltophilia Serine Proteases StmPr1 and StmPr2. Infect Immun 83:3825–37.

30. Karaba SM, White RC, Cianciotto NP (2013). Stenotrophomonas maltophilia encodes a type II protein secretion system that promotes detrimental effects on lung epithelial cells. Infect Immun 81:3210–9.

31. Berg G, Roskot N, Smalla K (1999). Genotypic and phenotypic relationships between clinical and environmental isolates of Stenotrophomonas maltophilia. J Clin Microbiol 37:3594–600.

32. Crossman LC, Gould VC, Dow JM, Vernikos GS, Okazaki A, Sebaihia M, et al. (2008). The complete genome, comparative and functional analysis of Stenotrophomonas maltophilia reveals an organism heavily shielded by drug resistance determinants. Genome Biol 9:R74.

33. Bi D, Liu L, Tai C, Deng Z, Rajakumar K, Ou HY (2013). SecReT4: a web-based bacterial type IV secretion system resource. Nucleic Acids Res 41:D660–5.

34. Schuster CF, Bertram R (2013). Toxin-antitoxin systems are ubiquitous and versatile modulators of prokaryotic cell fate. Fems Microbiology Letters 340:73–85.

35. El-Gebali S, Mistry J, Bateman A, Eddy SR, Luciani A, Potter SC, et al. (2019). The Pfam protein families database in 2019. Nucleic Acids Res 47:D427–D432.

36. Petersen TN, Brunak S, von Heijne G, Nielsen H (2011). SignalP 4.0: discriminating signal peptides from transmembrane regions. Nat Methods 8:785–6.

37. Sanchez MB (2015). Antibiotic resistance in the opportunistic pathogen Stenotrophomonas maltophilia. Front Microbiol 6:658.

38. Borgeaud S, Metzger LC, Scrignari T, Blokesch M (2015). The type VI secretion system of Vibrio cholerae fosters horizontal gene transfer. Science 347:63–7.

39. Sana TG, Flaugnatti N, Lugo KA, Lam LH, Jacobson A, Baylot V, et al. (2016). Salmonella Typhimurium utilizes a T6SS-mediated antibacterial weapon to establish in the host gut. Proc Natl Acad Sci U S A 113:E5044–51.

40. Anderson MC, Vonaesch P, Saffarian A, Marteyn BS, Sansonetti PJ (2017). Shigella sonnei Encodes a Functional T6SS Used for Interbacterial Competition and Niche Occupancy. Cell Host Microbe 21:769–776 e3.

41. Pompilio A, Crocetta V, Ghosh D, Chakrabarti M, Gherardi G, Vitali LA, et al. (2016). Stenotrophomonas maltophilia Phenotypic and Genotypic Diversity during a 10-year Colonization in the Lungs of a Cystic Fibrosis Patient. Front Microbiol 7:1551.

42. Brooke JS, Di Bonaventura G, Berg G, Martinez JL (2017). Editorial: A Multidisciplinary Look at Stenotrophomonas maltophilia: An Emerging Multi-Drug-Resistant Global Opportunistic Pathogen. Front Microbiol 8:1511.

43. Rouf R, Karaba SM, Dao J, Cianciotto NP (2011). Stenotrophomonas maltophilia strains replicate and persist in the murine lung, but to significantly different degrees. Microbiology 157:2133–42.

44. Di Bonaventura G, Pompilio A, Zappacosta R, Petrucci F, Fiscarelli E, Rossi C, et al. (2010). Role of excessive inflammatory response to Stenotrophomonas maltophilia lung infection in DBA/2 mice and implications for cystic fibrosis. Infect Immun 78:2466–76.

45. Alcoforado Diniz J, Liu YC, Coulthurst SJ (2015). Molecular weaponry: diverse effectors delivered by the Type VI secretion system. Cell Microbiol 17:1742–51.

46. Tang JY, Bullen NP, Ahmad S, Whitney JC (2018). Diverse NADase effector families mediate interbacterial antagonism via the type VI secretion system. J Biol Chem 293:1504– 1514.

47. Ting SY, Bosch DE, Mangiameli SM, Radey MC, Huang S, Park YJ, et al. (2018). Bifunctional Immunity Proteins Protect Bacteria against FtsZ-Targeting ADP-Ribosylating Toxins. Cell 175:1380–1392 e14.

48. LaCourse KD, Peterson SB, Kulasekara HD, Radey MC, Kim J, Mougous JD (2018). Conditional toxicity and synergy drive diversity among antibacterial effectors. Nat Microbiol 3:440–446.

49. Durand E, Cambillau C, Cascales E, Journet L (2014). VgrG, Tae, Tle, and beyond: the versatile arsenal of Type VI secretion effectors. Trends Microbiol 22:498–507.

50. Whitney JC, Quentin D, Sawai S, LeRoux M, Harding BN, Ledvina HE, et al. (2015). An interbacterial NAD(P)(+) glycohydrolase toxin requires elongation factor Tu for delivery to target cells. Cell 163:607–19.

51. Russell AB, Peterson SB, Mougous JD (2014). Type VI secretion system effectors: poisons with a purpose. Nat Rev Microbiol 12:137–48.

52. Si F, Li D, Cox SE, Sauls JT, Azizi O, Sou C, et al. (2017). Invariance of Initiation Mass and Predictability of Cell Size in Escherichia coli. Curr Biol 27:1278–1287.

53. da Silva AC, Ferro JA, Reinach FC, Farah CS, Furlan LR, Quaggio RB, et al. (2002). Comparison of the genomes of two Xanthomonas pathogens with differing host specificities. Nature 417:459–63.

54. Hayashi K, Morooka N, Yamamoto Y, Fujita K, Isono K, Choi S, et al. (2006). Highly accurate genome sequences of Escherichia coli K-12 strains MG1655 and W3110. Mol Syst Biol 2:2006 0007.

55. Hoang TT, Karkhoff-Schweizer RR, Kutchma AJ, Schweizer HP (1998). A broad-host-range Flp-FRT recombination system for site-specific excision of chromosomally-located DNA sequences: application for isolation of unmarked Pseudomonas aeruginosa mutants. Gene 212:77–86.

56. Welker E, Domfeh Y, Tyagi D, Sinha S, Fisher N (2015). Genetic Manipulation of Stenotrophomonas maltophilia. Curr Protoc Microbiol 37:6F 2 1–14.

57. Dykxhoorn DM, St Pierre R, Linn T (1996). A set of compatible tac promoter expression vectors. Gene 177:133–6.

58. Hachani A, Lossi NS, Filloux A (2013). A visual assay to monitor T6SS-mediated bacterial competition. J Vis Exp doi:10.3791/50103:e50103.

59. Vettiger A, Basler M (2016). Type VI Secretion System Substrates Are Transferred and Reused among Sister Cells. Cell 167:99–110 e12.

60. Ginestet C (2011). ggplot2: Elegant Graphics for Data Analysis. Journal of the Royal Statistical Society Series a-Statistics in Society 174:245–245.

61. Schindelin J, Arganda-Carreras I, Frise E, Kaynig V, Longair M, Pietzsch T, et al. (2012). Fiji: an open-source platform for biological-image analysis. Nat Methods 9:676–82.

62. Sievers F, Higgins DG (2018). Clustal Omega for making accurate alignments of many protein sequences. Protein Science 27:135–145.

63. Crooks GE, Hon G, Chandonia JM, Brenner SE (2004). WebLogo: A sequence logo generator. Genome Research 14:1188–1190.

64. Kumar S, Stecher G, Tamura K (2016). MEGA7: Molecular Evolutionary Genetics Analysis Version 7.0 for Bigger Datasets. Mol Biol Evol 33:1870–4.

65. El Yacoubi B, Brunings AM, Yuan Q, Shankar S, Gabriel DW (2007). In planta horizontal transfer of a major pathogenicity effector gene. Appl Environ Microbiol 73:1612–21.

